# Evaluating Multiomics Integration Architectures for Training With Structured Missingness

**DOI:** 10.1101/2025.09.10.672554

**Authors:** Simon Fisher, Jacob Bradley, George Lansdown, Owen Anderson, Russell Hung, James Lesh, Murray Cutforth, Ian Poole, Jeremy P. Voisey

## Abstract

Multimodal bioinformatics datasets are increasingly common in biomedical research, for tasks such as cancer subtyping and outcome prediction. It is feasible that data from a given patient, or even all data from a given institution, does not have coverage across all modalities; the available data is contingent on both the assay choice at the institution alongside technical aspects and associated drop-out. Consequently, algorithms for machine learning models must be tolerant to *structured* data missingness (or occurrence) for entire modalities when training. In this paper, we compare general strategies for training multimodal models in the context of structured modality missingness, employing suitable strategies for the stage of modality integration: *early* by concatenation of features, *intermediate* by max pooling of latent features, and *late* by aggregating model predictions probabilistically. We evaluate our strategies on a real-world bioinformatics dataset for the task of breast cancer subtyping, constructing a range of structured missingness scenarios. We highlight that, despite their inability to learn cross-modality interactions, late integration models outperform against early and intermediate integration strategies across a range of scenarios according to the level and nature of missingness. Logistic regression models, although simple, also outperform neural networks within the same settings. Fundamentally, we show that understanding the structure of missingness within a dataset is necessary when selecting a method of integration, and that simple models and approaches should not be dismissed when working with structured missingness.

## Introduction

Multimodal bioinformatics datasets, comprising biological data collected from multiple omics platforms, are becoming increasingly available thanks to the development of infrastructure for affordable acquisition of omics data (1–4). These datasets allow linkage and use of information over several layers of connected biology and can provide enhanced predictive signal across a variety of clinical prediction tasks.

When training (or applying) predictive models on multimodal bioinformatics data, there are challenges around integrating data modalities. Generally, integration strategies can be categorised according to three integration points: *early* integration, where modality features are concatenated prior to modelling; *intermediate* integration (also referred to as *middle* or *joint* integration), where modalities are modelled independently to produce latent modality representations which are then fused; and *late* integration, where the outputs of modality-specific models are combined (5). Several studies have developed intelligent integration strategies for multimodal (multiomic) data. For example, Lipkova et al. (6) presented the concept of “guided gradual fusion”, where more biologically related modalities (such as genomics and transcriptomics) may be integrated prior to those less related (e.g. imaging and genomics). Similarly, Shen et al. (7) developed the aggregate and map (AggMap) model, which leverages the pairwise correlation between multimodal datapoints in order to organise features into spatially correlated maps. Benkirane et al. (8) use a variant of intermediate integration, which they refer to as CustOmics. CustOmics uses autoencoders to fuse latent representations, using a custom loss function designed to reduce overfitting.

However, it is common to encounter data *missingness* in multimodal bioinformatics data. Missingness, also known as *incompleteness*, refers to missing observations of some features for some samples. In bioinformatics datasets, entire omics modalities tend to be missing rather than individual features, due to technical or cost constraints (9, 10). This is referred to as *structured* missingness. Structured data missingness presents a challenge for training machine learning models (11). There exist well established methods for handling feature-level missingness, including joint imputation and masking (5, 6), however these do not always transfer well to modality-wise structured missingness (12, 13).

A few works present multimodal integration strategies that handle structured data missingness in biomedicine. Cui et al. (14) classify glioma using an intermediate integration approach with missingness tolerance based on a combination of reconstruction loss and clustering-like classification loss. They train unimodal models to generate modality representations in comparable latent spaces, so that available modalities can be used to form a representation despite one modality being missing. Other authors have proposed methods such as cross-omics linked embedding with contrastive learning and self-attention (CLCLSA) (10), which are focused on prediction using partially missing data, but assume all data is available during training.

Outwith the bioinformatics domain, some studies have addressed training machine learning models in the presence of structured missingness. Lin and Hu (15) propose the use of modality-specific generative networks in order to synthetically reconstruct missing modalities. While applicable to bioinformatics data, this relies on robust single-modality representations which can only be learned from very large datasets- which are often lacking in biomedical research due to cost constraints. Ramazanova et al. (16) tackle missingness in non-biomedical data by learning missing modality tokens (MMTs), showing substantially higher performance than approaches which discard incomplete data. Lastly Lin and Hu (17), in a further paper argue against the use of single modality generative models, rather presenting a method of generating a fused embedding by learning cross-modality interactions.

To date, the impact of structured modality missingness on the choice of integration strategy has not been fully investigated for bioinformatics data. Therefore, the choice of whether to use early, intermediate, or late integration for a given biomedical task is generally guided by practitioner intuition rather than pre-existing empirical evidence obtained using different models on the same dataset. In this paper, we focus on the impact of missingness during training, aiming to expose the relationship between training set modality missingness and integration model performance across a *range* of architectures. For tractability we will not examine missingness at inference. Therefore we:

1. identify suitable mechanisms for integrating individual data modality representations in the presence of structured missingness, namely simple concatenation with modality drop-out, missingness-tolerant latent representation max-pooling, and missingness-tolerant independent encoding of modalities followed by probabilistic combination;
2. present experiments on the task of breast cancer subtyping using 3 data modalities in the publicly available Breast Invasive Carcinoma PanCancer Atlas from The Cancer Genome Atlas (TCGA), showing the robustness of different methods to structured missingness in the training set, with respect to both the overall level of missingness and degree of modality co-occurrence.

## Methods

Below we describe the (fully observed) dataset that we used, our method for simulating structured data missingness, and the three integration strategies that we evaluate.

### Datasets

We used data from the Breast Invasive Carcinoma PanCancer Atlas, obtained via cBioPortal (19), selecting the 3 data modalities described in Table 1. In order to maintain complete control over the presence of missingness in the training dataset, samples with missing features were dropped, leaving 981 samples with a full set of clinical, next generation RNA sequencing (RNA-Seq), and mutation data. Modality features were used in their original format other than for mutational data, where mutation counts were computed for each gene. Due to the very large experimental space formed we chose to perform a priori feature selection by filtering down to a subset of genes using knowledge from (18) as further described in supplementary, note 1. We show results from both feature selected models and all-feature models-but focus predominantly on feature-selected models.

**Table 1.** Input data modalities from Breast Invasive Carcinoma PanCancer Atlas used in our experiments, showing their raw size (“Unprocessed features”). “Selected features” are chosen according to the breast cancer biomarker panel outlined by (18), and used to train all modelsthough we also fit a subset of models on all features.

| Modality | Description | Raw Features | Selected Features |
| --- | --- | --- | --- |
| Clinical | Age, Sex, TNM<sup>1</sup> Staging | 5 | 5 |
| Transcriptomics | Gene-level count matrix | 20,531 | 66 |
| Genomics | Mutational count at gene loci | 18,987 | 68 |

For the target prediction task, we chose binary classification of the two dominant breast cancer subtypes within the cohort: invasive ductal (*n* = 780) versus invasive lobular (*n* = 201) breast cancer. This gave a total of 981 data samples, which were randomly allocated to train and test sets with an 80/20 split, stratified by subtype. The training set was used for data exploration and model development. The test set was held out and used to generate the results presented in this paper.

### Simulation of Missingness

We simulate *structured* missingness by modelling its inverse of “co-occurrence”, meaning the extent to which occurrence in one modality for a given sample *i* is correlated with occurrence in another modality. Letting *M*_*j*_ ∈ { 0, 1} ^*N*^ refer to the occurrence of modality *j* across *N* samples (*M*_*ji*_ = 1 if the modality *j* occurs for sample *i*), we define co-occurrence as the correlation of occurrence between any pair of modalities;

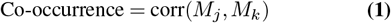

In our experiments we chose to have the same co-occurrence between every pair of modalities, with three settings of 0.9, 0.5, and 0.1. For each setting, we varied the proportion of missing samples equally for each modality from 0 to 0.7 in steps of 0.1, and from 0.7 to 0.95 in steps of 0.05.

For 3 modalities, there are 8 possible missingness patterns: [1, 1, 1], [1, 1, 0], [1, 0, 1], [0, 1, 1], [1, 0, 0], [0, 1, 0], [0, 0, 1], and [0, 0, 0]. To generate multiple random patterns of missingness yielding a specified degree of co-occurrence and missingness, we first calculated the number of each type of modality occurrence pattern required. Using ten different seeds for the random number generator, we then randomly chose training set samples to exhibit these.

Note that the test set was left complete, with no missing values.

### Integration Strategies

We chose representatives of three broad categories of approach to integration: early, intermediate, and late. These approaches are illustrated in Figure 1. The defining characteristics of these methodologies, and their implementation, are as follows.

**Fig. 1.**
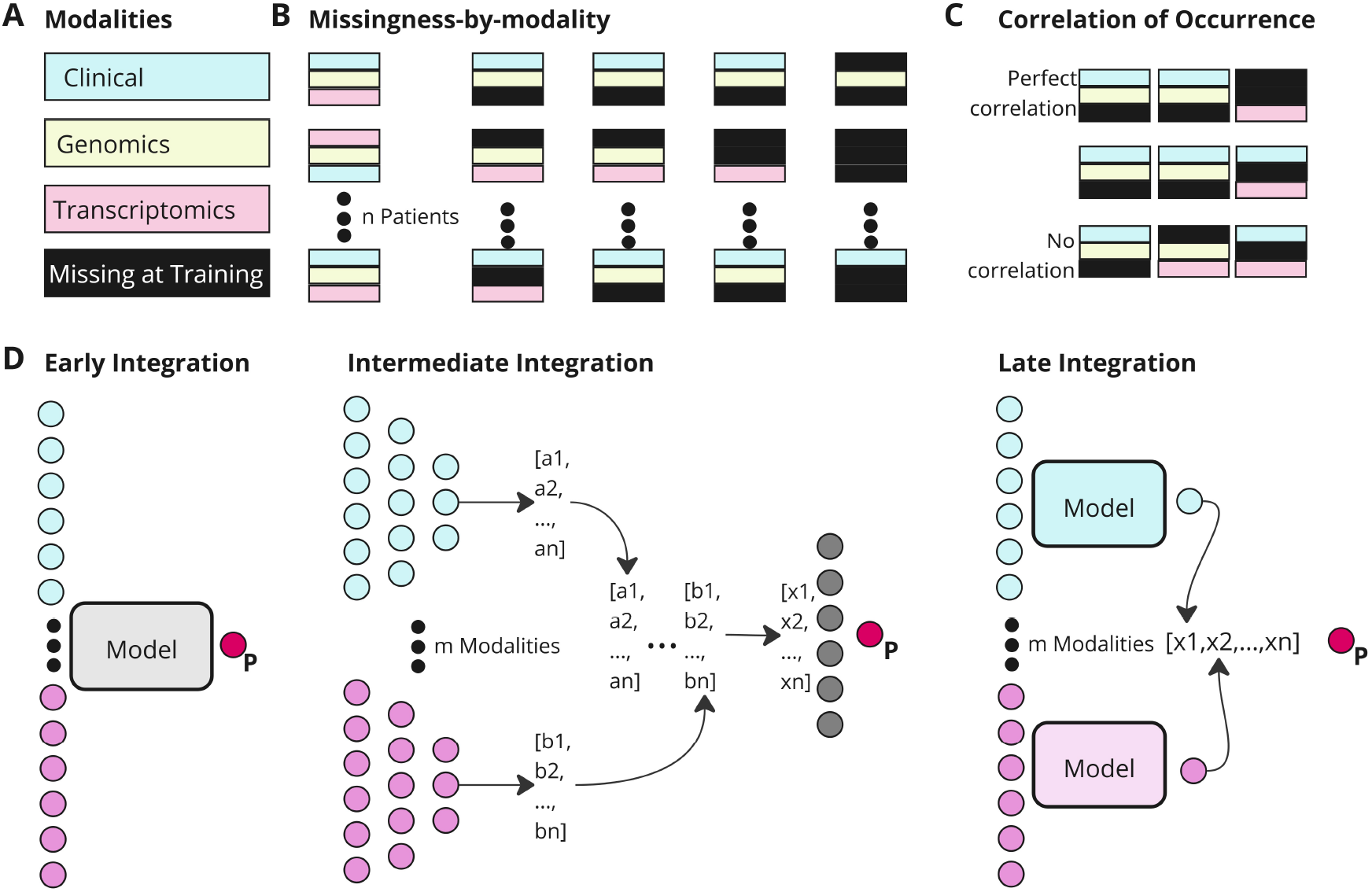
Overview of concepts. A) Clinical, Transcriptomics, and Genomics modalities are subjected to B) increasing degrees of missingness-by-modality. C) This missingness is assessed across a range of co-occurrence values. D) Visual outline of early, intermediate and late integration architectures. Nodes are coloured according to the modality of the features that are fed into that node; grey nodes receive features from multiple modalities. For intermediate integration, each modality produces a feature vector (in this case, ‘a’ and ‘b), which are fused into a combined vector (‘x’). “Model” indicates an object which constitutes either a fully connected network or a logistic regression classifier contingent on experiment.

#### M1:Early integration

We use ‘early integration’ to refer to any modelling strategy that directly concatenates feature vectors from input modalities. By extension, these models can only be trained using samples for which all modalities are observed and inference can only be performed if all modalities are available. As a representative of this strategy we used logistic regression, with *L*_2_ weight regularisation (weight selected by cross-validation performance and random search) or a shallow fully connected neural network with a single latent layer of dimension 16 trained for 25 epochs.

#### M2:Intermediate integration

We use ‘intermediate integration’ to refer to any approaches which encode each modality separately, and then combine these encoded representations in an intermediate latent space. Specifically, in our implementation each input modality is encoded using a shallow network, the outputs of which are equally sized latent representations which are then combined using an element-wise maximum pooling layer. Taking the maximum of each latent dimension across input modalities allows training (and inference) in the presence of modality-level missingness. The pooled latent representation is then passed through a final classification network. In the experiments presented below, we used two-layer encoder and single-layer classifier networks, with a single latent layer of dimension 16. We trained the model end-to-end for 25 epochs.

#### M3:Late integration

We use ‘late integration’ to refer to any modelling strategy that relies on training individual classifiers for each modality, and performing prediction using a method for ensembling output predictions. Since each classifier is trained on a single modality, missingness in other modalities does not impact the effective training set size. As a representative of this strategy, we employed logistic regression for each modality with *L*_2_ weight regularisation (weight selected by cross-validation performance and random search) or a shallow fully connected neural network with a single latent layer of dimension 16 trained for 25 epochs. We pool predictions probabilistically:

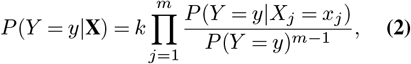

where *m* is the number of modalities, *Y* is the label for a given sample, *k* is a normalisation constant, and **X** = (*X*_1_, *X*_2_ … *X*_*m*_) are the sets of covariates from modalities 1, 2, …, *m*. The derivation of this expression is provided in supplementary, note 2.

For all methods described above, any hyperparameters were selected using 5-fold cross-validation. We present metrics of predictive performance applied to out-of-fold predictions in the training set. Models were refit to the entire training set before application to the test set.

## Results

In this section we present results on the held-out test set.

### Single Modality Classifiers

We trained unimodal classifiers to provide baselines, to allow us to gauge the benefit from multimodal classifiers. The results are shown in Figure 2.

**Fig. 2.**
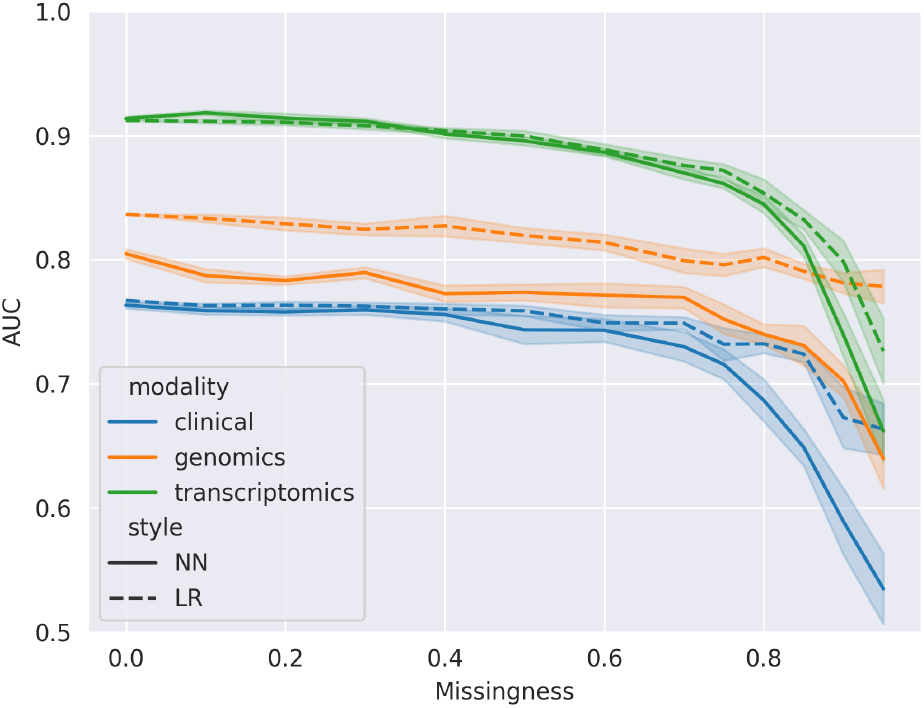
Performance of single modality classifiers across missingness proportions as measured by test-set AUC. Colour indicates single modality. Line style indicates the model type between neural-network (NN) or linear-regression (LR).

We observed that the transcriptomics classifier performed best across the full range of missingness values. It achieved a mean AUC of 0.912 on the fully observed dataset, with performance starting to drop at missingness values above 0.4 (i.e. only 60% or less of the training set available). In fact, all single modality classifiers (transcriptomics, genomics and clinical) show the same pattern of performance relative to missingness, with mean performance decreasing and performance variability increasing above 0.4. We also investigated the effect of missingness on the regularization of logistic regression models for single modality and early integration (Figure 3). Note that the C hyperparameter is the inverse of regularization strength. Transcriptomics was subject to the highest C hyperparameter (weakest regularisation) at low missingness. Regularization of genomics and clinical was stronger at low missingness, but progressively *weakened* with increasing missingness until it converged with transcriptomics at around 0.7 missingness. The early integration architecture using a logistic regression model followed a regularisation pattern similar to genomics and clinical.

**Fig. 3.**
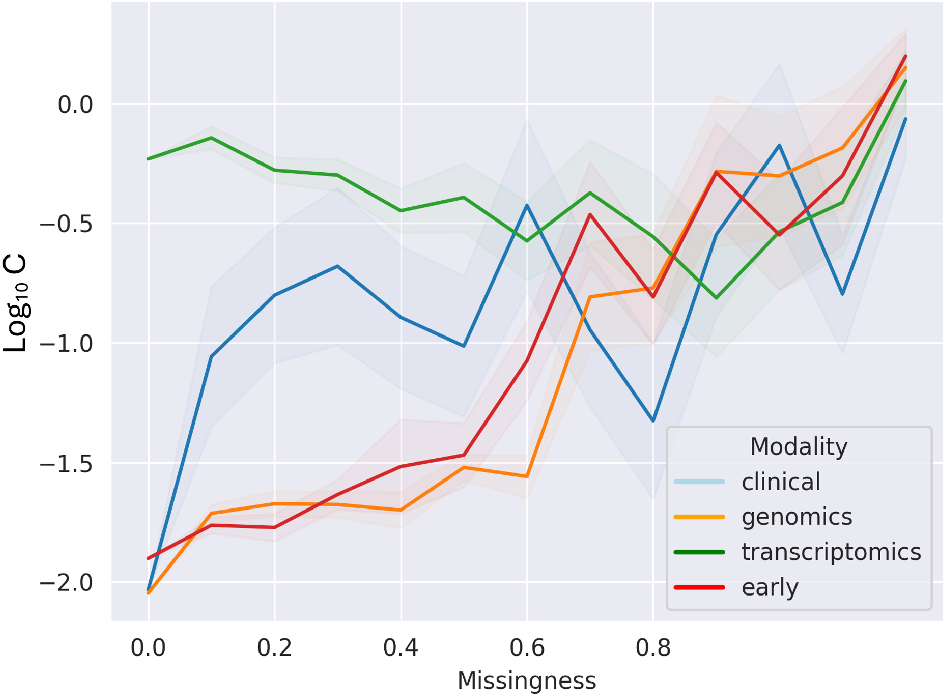
Optimal C parameter for logistic regression, as selected by 100-iteration gridsearch in five-fold cross validation. Early integration c-parameter also shown in red. Data shown by mean and standard error across 10 generated patterns of missingness.

### Multimodal Classifiers

The multimodal model results are shown in Table 2 and Figure 4. We show model performance for different integration types (colour), model types (line style), and panels (correlation in occurrence). These results were evaluated using the holdout test set using ten patterns of missingness. Firstly, at 0 missingness the late integration architecture was consistently the best performing approach across all correlations of occurrence (0.9, 0.5, and 0.1). It continued to be the best performing across increasing missingess, showing both the highest starting performance AUCs, 0.938 (LR) 0.930 (NN), and the greatest resistance to the injection of missingness, only decreasing to 0.875 (LR) and 0.856 (NN) at 0.8 missingness. By contrast, early integration architectures, whilst performing well at 0 missingness, AUCs were 0.931 (LR) and 0.926 (NN), were highly intolerant to increasing levels of missingness. This was particularly prominent in a scenario of low correlation in modality occurrence, where we observed a reduction to 0.772 (LR) and 0.704 (NN) at 0.8 missingness. However, the introduction of mean imputation to logistic regression models effectively reversed the vulnerability of early integration to missingness (Figure 5). We also supported our headline findings by examining integration models without using a priori feature selection and observed worsened performance across all missingness parameters tested- which was particularly apparent with neural network models (Figure 6)

**Table 2.**
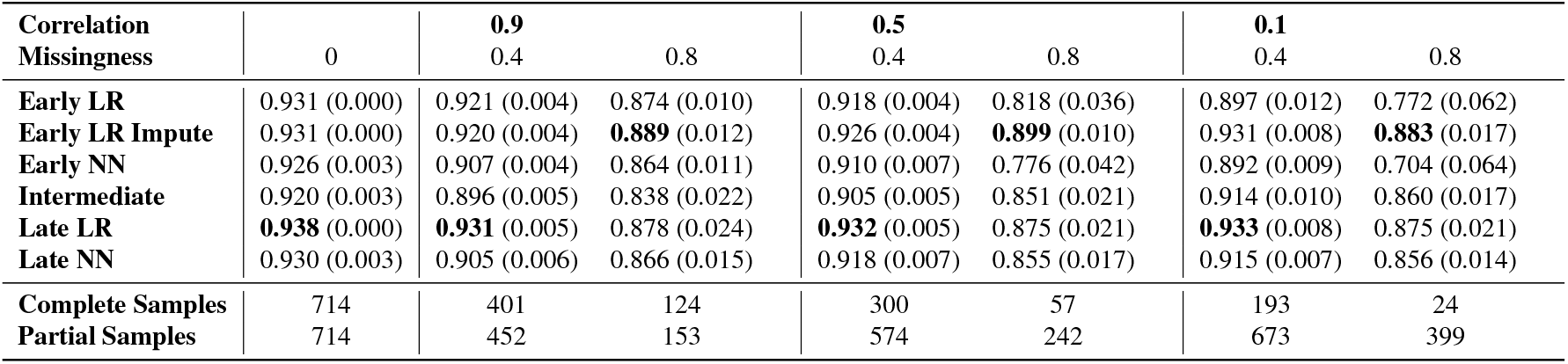
Model performance (AUC) of models trained under different selected correlation in occurrence (0.9, 0.5, 0.1) and missingness proportions (0, 0.4, 0.8) within those correlations. Models evaluated on the holdout test set which comprises fully observed data. Number observed training samples by both complete (all modalities) and partial (one modality). Variability across the 10 generated patterns of missingness assessed is represented by 95% confidence interval shown in parenthesises.

**Fig. 4.**
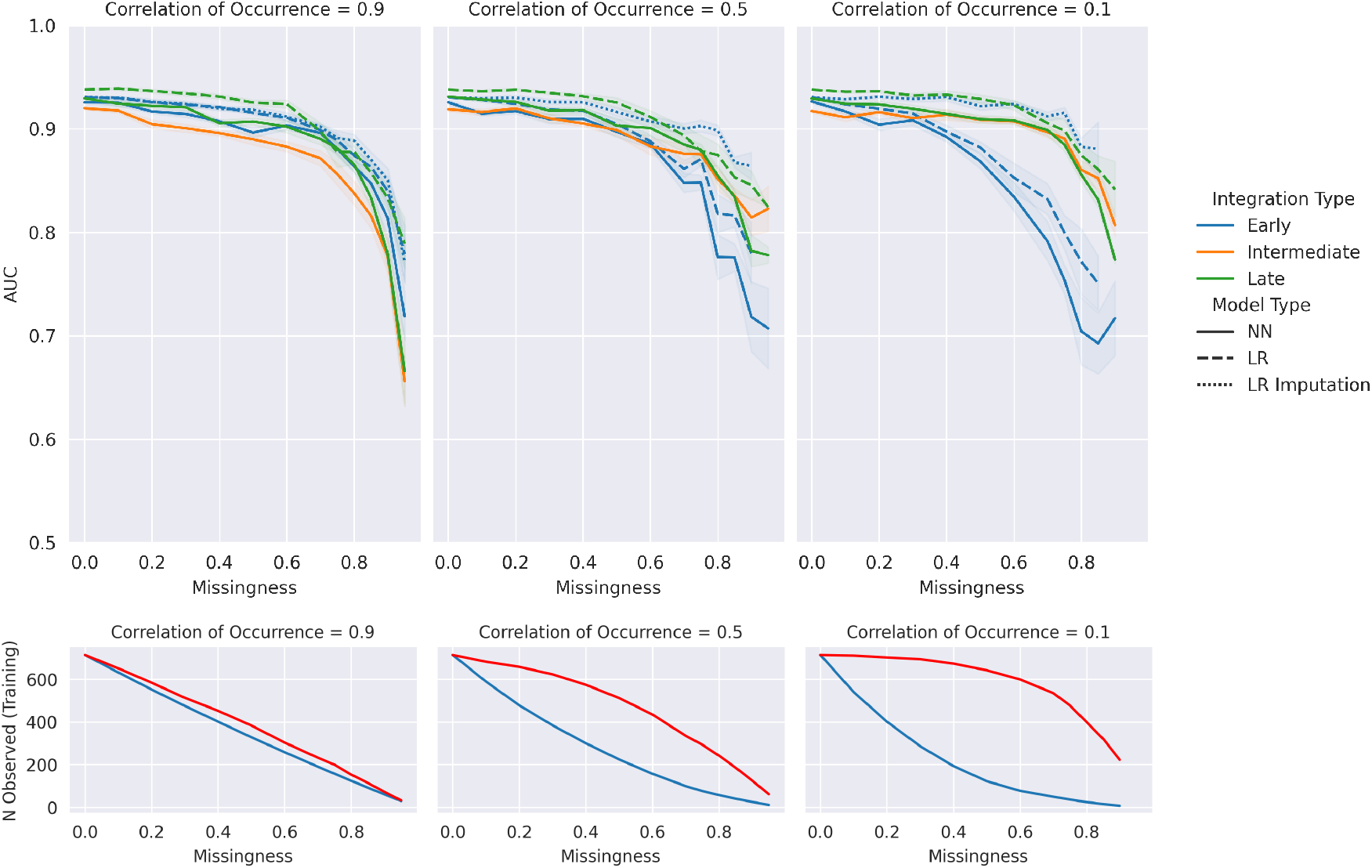
Top: Performance of multimodal classifiers by AUC (Early (blue), Intermediate (orange), Late (green), by model-type (Neural network (filled), logistic regression (dashed)) across missingness proportions and correlation of occurrence on the holdout test set. Data shown by mean and standard error across 10 generated patterns of missingness. Bottom: Training set sample size (n) where a sample has at least one available modalities (red) or all available modalities (blue).

**Fig. 5.**
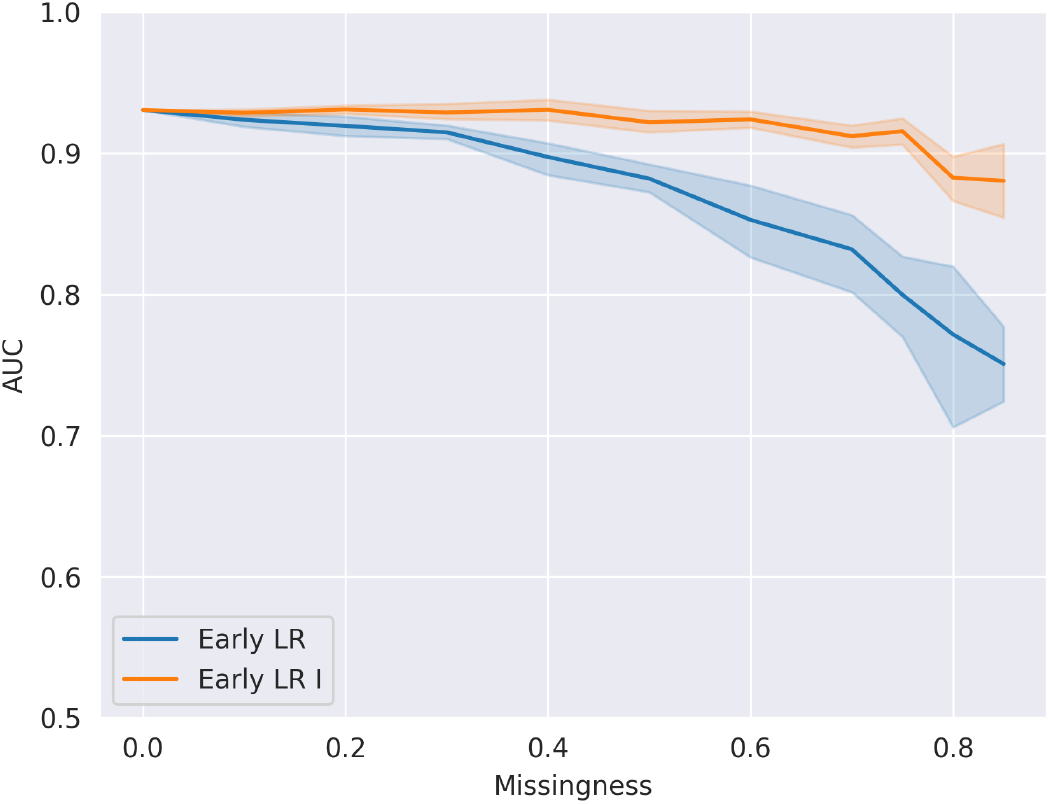
Performance of Early logistic regression classifiers with (orange) or without (blue) mean imputation of features in holdout test set at 0.1 correlation of occurrence. Data shown by mean and standard error across 10 generated patterns of missingness.

**Fig. 6.**
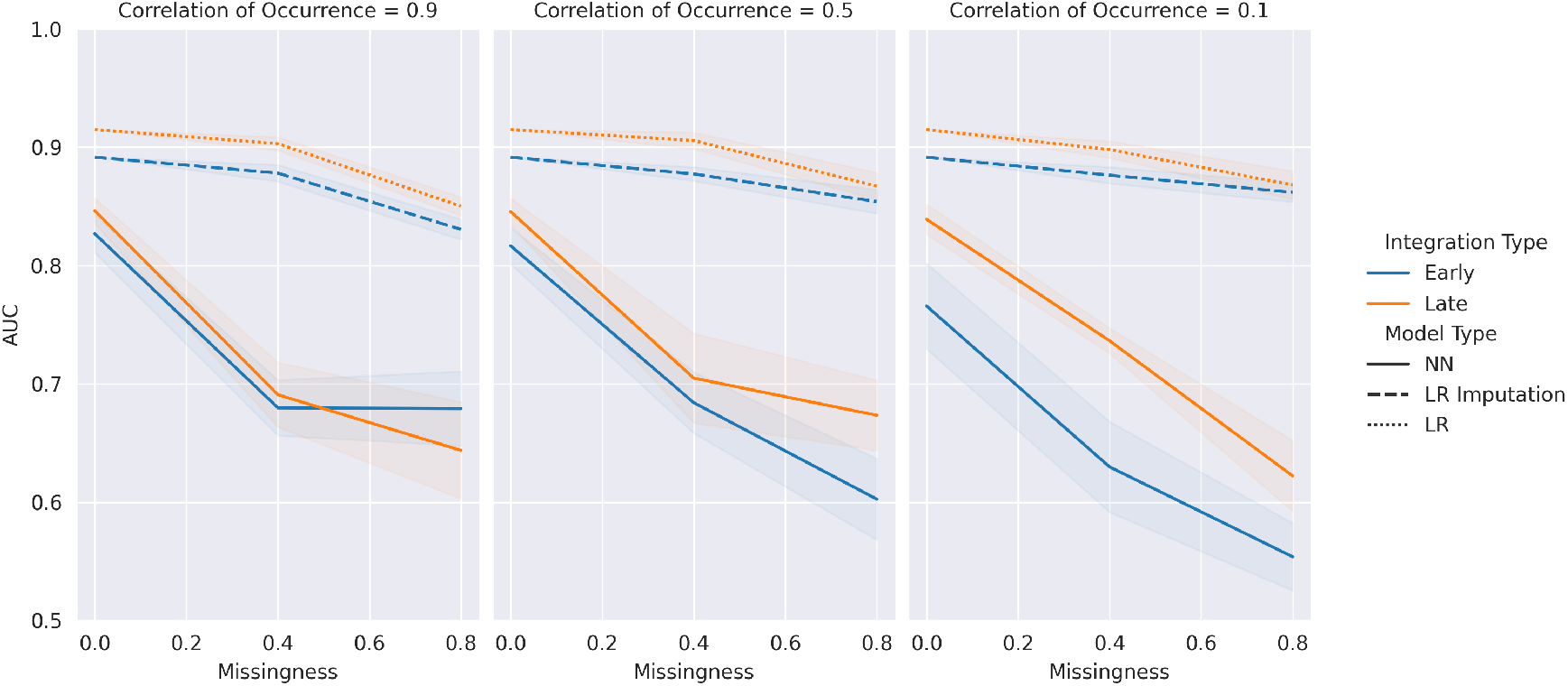
Performance of Early (blue) or Late (orange) architectures where no feature selection was performed for transcriptomic or genomic modalities. Models fitted to data of limited missingness values of 0, 0.4 and 0.8 only due to fitting time. Data shown by mean and standard error across 10 generated patterns of missingness.

We note that logistic regression models used with both early and late architectures consistently outperformed their neural network counterparts by margins of between 0.01 and 0.15 AUC contingent on missingness and correlation of occurrence.

To accompany these findings we also calculated the availability of samples for the various models from the training cohort (Figure 4). Where early integration without imputation requires samples with complete modalities to be used in training, intermediate, late, and early with imputation only require at least one modality to train with that sample. Thus, utility of training data is maximized for intermediate, late, and early with imputation integration.

Intermediate integration was the poorest performing architecture at high correlation of occurrence (0.9) across differential levels of missingness. Interestingly however, it was the only method which performed better at low levels of correlation of occurrence than high (Figure 7). At this a low correlation of occurrence (0.1) it becomes competitive with the early and late approaches as seen in Figure 4.

**Fig. 7.**
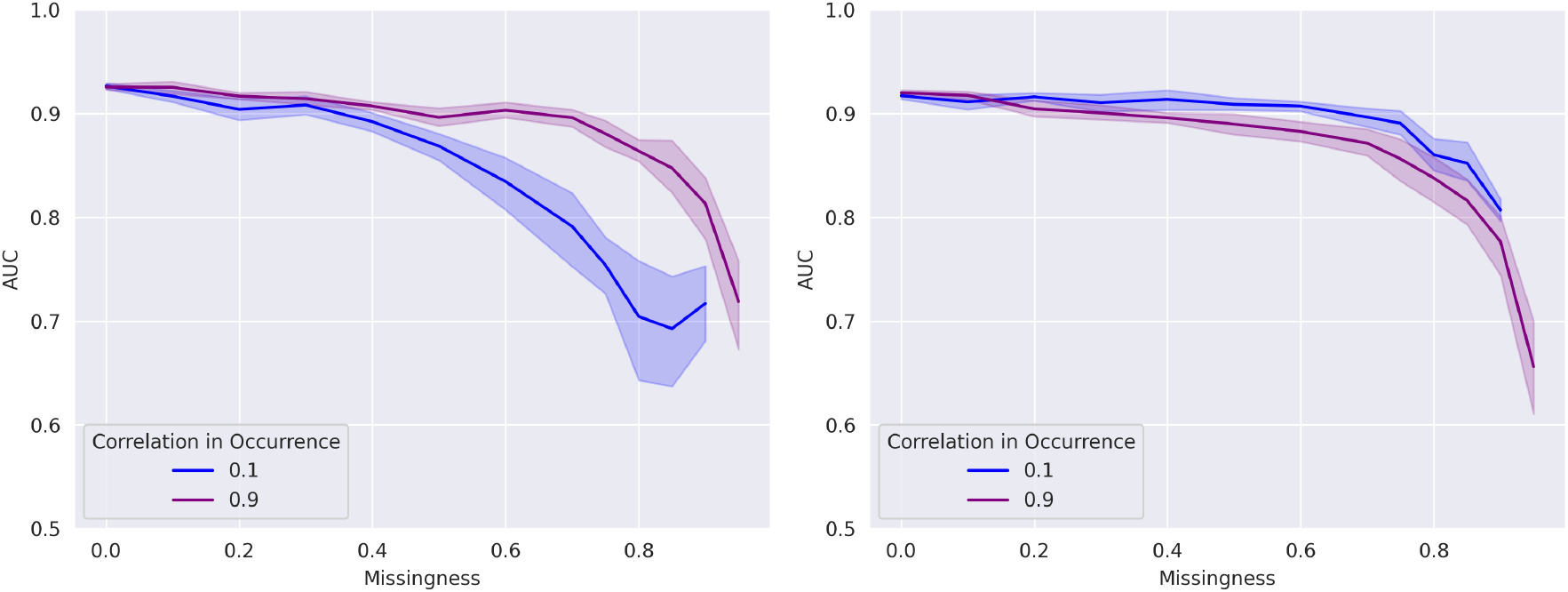
Model performance (AUC) of specific early and intermediate neural network models across missingness and the extremes of correlation in occurrence in the holdout test set. Data shown by mean and standard error across 10 generated patterns of missingness.

## Discussion

In this paper, we examined a range of modality-level missingness proportions which were further stratified by cooccurrence of modalities. On one extreme, we created a high co-occurrence scenario where modalities were generally missing or occurring together (a reasonable scenario in the context of omics modalities), but also investigated low co-occurrence where missingness was highly random. As expected, the performance of all models and approaches decreased as missingness increased- attributed to the fundamental reduction in the number of samples available to train models. However, there were variations in the mangitude of this effect across experiments. The findings of our study indicate that across the three primary integration architectures, early, intermediate, and late, late integration is the most consistently high performer across scenarios within our clinical, genomics and transcriptomics dataset. Additionally, we observed that logistic regression based classifiers tend to outperform their neural network counterparts across all scenarios. Finally, we note that even simple approaches to modality imputation significantly rescue performance even with high degrees of missingness.

The central dogma of biology — the broadly one-directional translation from DNA to physiological effect — means that there is physical linkage between omics modalities. For example, overexpression of a mutated protein may have different disease implications to overexpression of a wild-type protein. It is reasonable, therefore to assume that making this linkage of modalities available to a model could be beneficial to a task, as it allows cross-modality interactions to be learnt. Late integration, because it segregates modalities during modelling, can be regarded as the architecture which is least applicable to the sharing of information across the separate modalities. Two groups in their reviews therefore effectively dismiss it on the basis of this assumption (20, 21).

Here, we clearly and consistently show that despite violating the assumption of modality linkage, simple late integration of modalities with probabilistic pooling is the best performing approach. The failure, in our study, of early logistic regression and neural network models to out-compete late integration, even where imputation rescues performance, indicates that in our dataset (and perhaps more widely for multimodal omics) cross-modality interactions are not necessary for model performance. This is an observation previously made by (22) where they do not show a difference between simple support vector regression and neural networks consisting of both MLP and their state-of-the-art model in the two datasets they assessed. The inability of neural networks to outperform logistic regression using early integration with no missingness even suggests that intra-modality interactions may not be essential. We acknowledge that this is contradictory to (23), who via their InterSHAP network present the utility of cross-modal interactions between RNA and protein for single cell classification. However, we also note that simple classifiers (non-neural networks) were not tested in their experiments and thus cannot be benchmarked against.

It should be noted that in our experiments with multiple variations of datasets (co-occurrence, missingness, random patterns with different seeds) we did not individually optimise neural network design and training, which is something a practitioner working with a single dataset would do. Even so, simple classifiers may set a high baseline, which may not necessarily be surpassed. Despite this, comprehensive published material on multiomic integration using simple classifiers is challenging to find. We hypothesise that researchers tend to favour complex, deep, generative or graph-based networks-perhaps because they anticipate the need for complex modelling spaces in reflection of the complex underlying biology from which the data was derived. To be clear, we are not denying the existence of cross-modality interactions, as these fundamentally underpin biology and are empirically demonstrated. Rather, we are making the case that it is very difficult for highly-parameterised models to separate these from noise within datasets of modest or realistic size. Complex models, while theoretically capable of learning true interactions, are also at risk of finding spurious ones; i.e. over-fitting; this is further supported by our finding that *a priori* feature selection significantly boosts performance in the designated task versus using all available features.

Alongside cross-modality feature space not being necessary, we believe the higher performance of late integration versus its counterparts (particularly as missingness increases) is explained by two technical reasons. Firstly, late integration allows for the maximisation of data utility compared to early (without imputation). The importance of larger sample sizes when training models is well-known (24). The degradation in performance as missingness increases and sample size decreases supports this. Early integration without imputation discards any samples that are incomplete, whilst late integration makes use of any samples having at least one modality. Even in a situation of imputation, however, late integration still outperforms early, albeit by a much smaller margin. Furthermore, analysis of the regularisation hyperparameter for logistic regression models indicated another consideration. Single modality models that are used in late integration are better able to select appropriate hyperparameters for their modalities than when these modalities are fused or directly concatenated during early integration. This allows for superior optimisation of models, indicated by the clear separation in C between single modalities. The analysis of the regularisation hyperparameter does raise a question however. At zero missingness, modalities with poorer individual performance (clinical and genomics) tend to have stronger regularisation, which seems intuitive. The strength of this regularisation drops, however, as missingness increases. We would expect the opposite: that as sample sizes decrease stronger regularisation is required. This may indicate a failure of crossvalidation to find the optimal hyperparameter as datasets decrease in size and warrants further investigation.

In this study we have focussed on missingness at training; modalities may also be missing at inference. While it is evidently possible to use some models to make predictions with missing modalities, there is an alternative approach: a suite of models could be trained for each combination of potential modalities. At inference a model that has been optimised for the available modalities of a sample could then be used. We suggest that this approach would be preferable to a single model that can deal with missing modalities, but has not been optimised for those available in the test sample. While this may present difficulties with a large number of combinations, practical considerations could limit these to a manageable number. In addition, a machine learning pipeline could readily be used to generate this suite of models. When evaluating the approaches, we have assumed that three modalities: Clinical, Transcriptomics, and Genomics are present and have optimised models accordingly. This is equivalent to ignoring additional hypothetical modalities that were present in the training data, but not available at inference.

In summarisation, we suggest our findings indicate that maximisation of data availability on training is paramount, and that simple approaches should be continue to be properly investigated.

We acknowledge this study focuses on a single task with three modalities of data- two being omics modalities. Breast cancer sub-typing clearly displays a high level of predictive signal at baseline, and thus the conclusions we draw here are applicable to datasets where the researcher might expect a strong biological signal. However, we encourage readers to consider the implications regardless of signal strength. Additionally, simple machine learning architectures cope with tabular data whilst they are intolerant to more complex modalities such as imaging or text data. Where encoders or transformations are required (such as would be the case for a wholeslide-image modality or a radiology report) end to end models would need to be deep-learning in nature. This is especially pertinent where those encoders were expected to be trained in the context of a label. However, given this limitation, we encourage readers to consider blended approaches and use simple models where possible. Missingness was introduced synthetically so that modalities are missing completely at random (MCAR), and their missingness is not informative. In reality, the missingness of a modality may be missing at random (MAR) or missing not at random (MNAR) and may be informative. Finally, whilst we elected to focus on missingness at training in this study; modalities may also be missing at inference. In fact, it is reasonable to assume that missingness patterns in inference data would mirror those in training data. We emphasise that the frameworks here are tolerant to missingness at inference but providing this dimension to our study would have significantly complicated our ability to report findings. Researchers should maintain awareness within the conclusions of this study that they relate only to fully observed test sets.

## Conclusion

In this paper we evaluated a number of approaches for integrating multimodal datasets across a range of missingness and co-occurrence scenarios. This evaluation was carried out using a real-world dataset (from the Breast Invasive Carcinoma PanCancer Atlas) on a genuine task: cancer subtyping. Despite the inability of late integration to allow cross-modality interaction, we showed that it performed consistently well. Further logistic regression which does not allow for intra-modality interactions also performed well. These findings suggest that researchers should maximise their training data availability and aim for simple models where possible

## Acknowledgments

The results shown here are in whole or part based upon data generated by the TCGA Research Network: https://www.cancer.gov/tcga.

## Supplementary Note 1: Gene list

PIK3CA, RUNX1, CDH1, TP53, TBX3, PTEN, FOXA1, MAP3K1, GATA3, AKT1, NBL1, KMT2C, DCTD, RB1, SF3B1, CBFB, ARHGAP35, OR9A2, NCOA3, RBMX, MAP2K4, TROVE2, NADK, CASP8, CTSS, ACTL6B, LGALS1, KRAS, KCNN3, FBXW7, LRIG2, PIK3R1, PARP4, ZNF28, HLA-DRB1, ERBB2, ZMYM3, RAB42, CTCF, ATAD2, CDKN1B, GRIA2, NCOR1, HRNR, GPRIN2, PAX2, ACTG1, AQP12A, PIK3C3, MYB, IRS4, TBL1XR1, RPGR, CCNI, ARID1A, CD3EAP, ADAMTS6, OR2D2, TMEM199, MST1, RHBG, ZFP36L1, TCP11, CASZ1, GAL3ST1, FRMPD2, GPS2, and ZNF362.

## Supplementary Note 2: Derivation of Late Integration

Naivë Bayes (assumes **conditional independence** for *X*_*i*_) states:

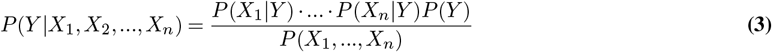

In the usual application of Naivë Bayes *X*_*i*_ is an individual feature. Note that in our case *X*_*i*_ is a modality, not a single feature. Hence we are assuming conditional independence of **modalities**:

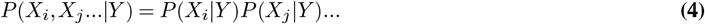

Bayes theorem states:

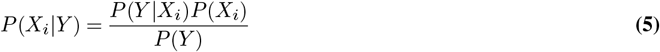

Substituting into the Naivë Bayes formula, for *X*_*i*_:

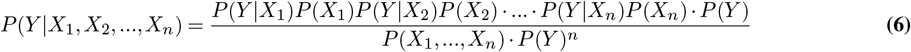

This can be rearranged to:

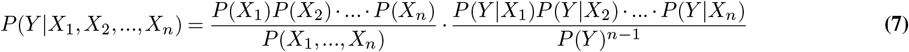

In addition to the assumption of conditional independence, we need to make the stronger assumption of independence of modalities:

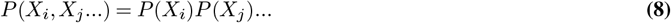

Using this assumption, the first fraction in the expression cancels out:

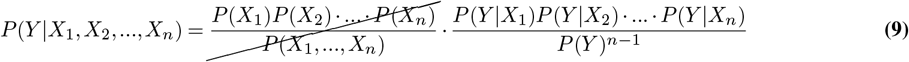

Thus, our equation can be written as:

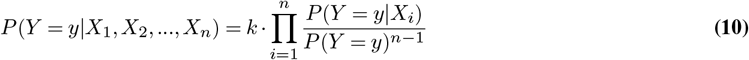

where k is a normalisation constant, to ensure the value of the probability falls within 0 and 1, and is necessary due to the assumptions and approximations made.

